# BindingGYM: A Large-Scale Mutational Dataset Toward Deciphering Protein-Protein Interactions

**DOI:** 10.1101/2024.12.03.626712

**Authors:** Wei Lu, Jixian Zhang, Ming Gu, Shuangjia Zheng

**Affiliations:** Aureka Biotechnologies

## Abstract

Protein-protein interactions are crucial for drug discovery and understanding biological mechanisms. Despite significant advances in predicting the structures of protein complexes, led by AlphaFold3, determining the strength of these interactions accurately remains a challenge. Traditional low-throughput experimental methods do not generate sufficient data for comprehensive benchmarking or training deep learning models. Deep mutational scanning (DMS) experiments provide rich, high-throughput data; however, they are often used incompletely, neglecting to consider the binding partners, and on a per-study basis without assessing the generalization capabilities of fine-tuned models across different assays. To address these limitations, we collected over ten million raw DMS data points and refined them to half a million high-quality points from twenty-five assays, focusing on protein-protein interactions. We intentionally excluded non-PPI DMS data pertaining to intrinsic protein properties, such as fluorescence or catalytic activity. Our dataset meticulously pairs binding energies with the *sequences* and *structures of all interacting partners* using a comprehensive pipeline, recognizing that interactions inherently involve at least two proteins. This curated dataset serves as a foundation for benchmarking and training the next generation of deep learning models focused on protein-protein interactions, thereby opening the door to a plethora of high-impact applications including understanding cellular networks and advancing drug target discovery and development.

## 1 Introduction

Protein-protein interactions (PPI) represent a vital component of the cellular language, mediating communication within and between cells [1, 2]. The strength of these interactions is commonly measured experimentally as the binding free energy, denoted Δ*G*, or referred to as binding affinity. In antibody drug discovery, a primary optimization goal is to enhance affinity towards desired targets (affinity maturation) while reducing affinity towards non-desired targets. For example, a broad-spectrum neutralizing antibody drug should bind strongly to prevalent variants of COVID-19 virus proteins to prevent immune escape, yet it should not be polyreactive [3, 4].

Despite its importance, progress in predicting binding affinity is limited by the scarcity of publicly available data from low-throughput biophysical experiments. While high-throughput techniques such as Yeast Two-Hybrid and affinity purification-mass spectrometry (AP-MS) provide extensive binary PPI data, these methods only indicate whether proteins bind, lacking details on the strength of the interactions [5, 6]. Additionally, due to the nature of AP-MS, binary PPI data may include pairs that do not directly interact but are connected through a third protein [7]. Another high-throughput method, deep mutational scanning (DMS), maps genotype to phenotype by combining screening techniques such as fluorescence-activated cell sorting (FACS) with next-generation sequencing (NGS) [8, 9]. This method can produce quantitative fitness scores for millions of mutant proteins sequenced by NGS. While DMS has been used to study a wide range of protein properties, including stability and fluorescence, our dataset exclusively includes studies examining protein-protein interactions, where the fitness score correlates with PPI binding affinity. Unlike previous protein-related datasets that only include information about the mutated protein [10, 11, 12, 13], our dataset explicitly incorporates all interacting proteins. The binding affinity between protein A and B differs from that between A and C, and proteins rarely interact with only a single partner. Predicting the binding affinity based on a single protein, such as protein A alone, is often not meaningful; models trained on such incomplete data fail to capture residue-level interactions and struggle to generalize. Including information about all interacting partners enables the training of a more generalizable PPI model across different assays. Furthermore, in some studies, the binding affinities between a mutant protein and multiple distinct interaction partners are individually measured [14]. Previously, this situation could result in multiple, potentially conflicting affinity data points for each mutant. Now, with complete partner information available, these data points become invaluable, enabling models to discern the intricate details of interaction specificity.

Currently, DMS results are predominantly used by sequence-based methods, whereas structure-based methods, traditionally preferred for estimating protein-protein binding energy, rarely leverage such data. To enable structure-based models to also benefit from DMS results and to facilitate the development of structure-based deep learning models, thus ensuring a level playing field between structure and sequence-based approaches, we map the wild-type sequences documented in source papers to their corresponding crystallized complex structures in the Protein Data Bank [15] through a comprehensive pipeline. For sequences that do not precisely match with the sequences in the crystal structures, we employ homology modeling using BioPython and OpenMM [16, 17]. Additionally, to enable our dataset to support baseline models that require both the wild-type and mutant structures, we use FoldX [18, 19] to generate the complex structure for each mutant.

In BindingGYM, we have assembled the largest collection of DMS-based PPI data available, gathering over ten million raw DMS data points and refining them into half a million high-quality data points from twenty research papers. Each entry includes complete data: the binding energy score, sequences of all interacting proteins, and the structure of the entire complex. This completeness allows our dataset to support a broad range of modeling approaches, including both sequence-based and structure-based methods. Additionally, we have introduced two novel and practically important data-splitting strategies: ’Central vs. Extremes Split’ and ’Inter-Assay Split’. The first strategy trains models on entries with middle-range binding energies, from the 10th to the 90th percentile, and evaluates them on the extremes, while the second strategy utilizes data from multiple assays to train models that predict outcomes in unseen assays. Similar to how the ImageNet dataset [20] has been foundational in advancing deep learning models for computer vision, and the high-throughput SELEX dataset [21, 22] in training AlphaFold3 for protein-DNA structure prediction [23], BindingGYM is poised to drive significant advancements in the field of protein-protein interactions. All scripts and data are freely accessible at https://anonymous.4open.science/r/BindingGYM-602D/.

## 2 Related Work

### Protein self properties

Several datasets are available to evaluate model performance on a range of protein properties, including secondary and tertiary structures, catalytic activity, stability, and expression. Early work, such as TAPE [11], focused primarily on structural properties, designing tasks for secondary structure prediction, contact prediction, and overall structure, as well as two engineering properties: fluorescence and stability. Meanwhile, FLIP [12] introduced multiple splitting schemes for evaluating protein engineering properties but limited its assessments to results from three assays, focusing solely on sequence-based models.

ProteinGYM [13] provides a comprehensive collection of DMS and clinical variants data. It standardizes measurements under a single metric, the fitness score, which is effective for zero-shot evaluation of models across a broad spectrum of protein properties. However, its fine-tuning capabilities are confined to individual assays, lacking generalization to test cross-assay performance for fine-tuned models. Importantly, while ProteinGYM includes binding-related assay results, it only provides data for the protein undergoing mutation, neglecting its interacting partners. Additionally, critical data are often missing; for instance, although [14] conducted screenings for the KRAS protein against seven different proteins, ProteinGYM includes results for only one. Similarly, while [24, 25] contain multiple mutations, ProteinGYM only includes data for single mutations.

### Low throughput quantitative PPI dataset

Due to the importance of protein-protein interactions, many datasets have been carefully curated, collecting binding affinity measurements from hundreds of papers [26, 27, 28, 29]. However, constrained by the low throughput of conventional biophysical methods, the most comprehensive dataset, SKEMPI [30], comprises only 7,085 data points and is heavily biased toward Alanine substitutions. BindingGYM can be considered the next-generation SKEMPI dataset; like SKEMPI, it includes the full complex structure with binding score but provides orders of magnitude more data, enabling the training of advanced deep learning models, as listed in Table 1 It is important to note that the DMS score does not directly equal Δ*G* but correlates with it. Consequently, the absolute values of DMS scores are not comparable across different assays. To address this, proper grouping of training samples is crucial, and methods such as learning-to-rank techniques [31, 32, 33] should be employed.

**Table 1:** Dataset Comparison: BindingGYM offers the most extensive collection of quantitative protein-protein interaction data currently available. Additionally, each data is meticulously paired with its corresponding protein complex structure, facilitating comparisons between sequence-based and structure-based methods. Detailed definitions of each column are provided in SI. HT: High-throughput Assay, C-Structure: Complex structure, PGYM: ProteinGYM, PW: ProtienWorkshop

| Dataset | # of data | HT | C-Structure Available | Quantitative | ML ready | Multichain support | Design Usecase |
| --- | --- | --- | --- | --- | --- | --- | --- |
| BindingGYM | 10M | ✓ | ✓ | ✓ | ✓ | ✓ | protein-protein interaction |
| SKEMPI [30] | 7K | ✗ | ✓ | ✓ | ✗ | ✓ | protein-protein interaction |
| STRING [35] | 300M | ✓ | ✗ | ✗ | ✗ | ✗ | protein network |
| PGYM [13] | 2.7M | ✓ | ✗ | ✓ | ✓ | ✗ | general protein fitness |
| FLIP [12] | 320K | ✓ | ✗ | ✓ | ✓ | ✗ | general protein fitness |
| TAPE [11] | 100K | ✗ | ✗ | ✓ | ✓ | ✗ | protein representation learning |
| PW [38] | 2.3M | ✓ | ✗ | ✓ | ✓ | ✗ | protein representation learning |
| FLAb [39] | 10K | ✗ | ✗ | ✓ | ✓ | ✗ | therapeutic antibody design |

### High throughput binary PPI dataset

Binary PPI provides protein network information at a proteome scale, which is invaluable for identifying key proteins underlying diseases or catalytic pathways. High-throughput methods yield binary (bind/non-bind) data for various model systems, with databases such as STRING and BioGRID containing millions of binary PPI entries [34, 35]. However, the scores in these databases reflect confidence levels in the existence of interactions rather than their strength. Consequently, models trained on this binary data tend to focus on protein-level evolution instead of residue-level physics, which are crucial for generalizing predictions of mutational effects [36]. While datasets like those in [37] exist, BindingGYM distinguishes itself by specifically focusing on mutational effects, rather than broad proteomic network predictions.

## 3 The BindingGYM dataset

### 3.1 Data collection

We collected over ten million DMS data points for 41 unique protein-protein complexes from twenty research papers, specifically focusing on protein-protein interactions. The complete list can be found in the supplementary material. For each paper, we manually traced the raw NGS data whenever possible, checked the data distribution to ensure consistency with the conclusions of the original papers, and corrected any errors introduced during data processing. For example, we identified and corrected misaligned entries in the processed data provided by Heredia et al. [40] by consulting the original article and its raw data.

Unlike ProteinGYM [13], which uses UniProt [41] sequences as references, we ensure our reference sequences are those actually used in the experiments by verifying against the original articles, appendices, and source codes. This required significant manual effort to identify the actual plasmids and reference sequences used for screening experiments.

In BindingGYM, we find the closest complex structures in the PDB for articles that contain DMS data but lack complex structures, facilitating the application of structure-based methods. When discrepancies arose between PDB structures and actual reference sequences, we employed homology modeling [17, 16] to align the structures with the reference sequences. To support baseline models that require both wild-type and mutant structures, we used FoldX [18, 19] to generate the complex structure for each mutant. The FLIP dataset [12] comprises data from three papers, including one on binding interactions that we have incorporated into our study. Three assays [42, 43, 44] classified as binding-related in ProteinGYM are not actually related to binding and were therefore excluded. Additionally, we noted dataset coverage discrepancies: the original paper [14] reports mutant proteins against seven targets, whereas ProteinGYM includes only one, and covers only single mutations while multiple mutations are documented [24, 25]. We also developed a refined dataset with 508,962 data points, by setting higher NGS count thresholds and applying additional filters, as detailed in the supplementary materials, to facilitate model benchmarking.

### 3.2 Data splits

As noted in [12, 13], the choice of data splitting scheme is crucial for a fair comparison between models. Random splits often overestimate model generalization because realistic objectives typically involve designing stronger-binding mutants, extrapolating to unexplored fitness landscapes, or applying models to new proteins. Consequently, alternative splitting schemes are necessary to ensure more accurate evaluations. As depicted in Figure 1, we have implemented five distinct data splitting schemes. The first, ’Conti Split’, separates continuous blocks in sequence space for both training and testing. The second, ’Mod Split’, assigns every n-th residue in sequence space to the test set. The third, ’Rand Split’, represents the commonly used random split. These initial three schemes are also used in ProteinGYM.

**Figure 1:**
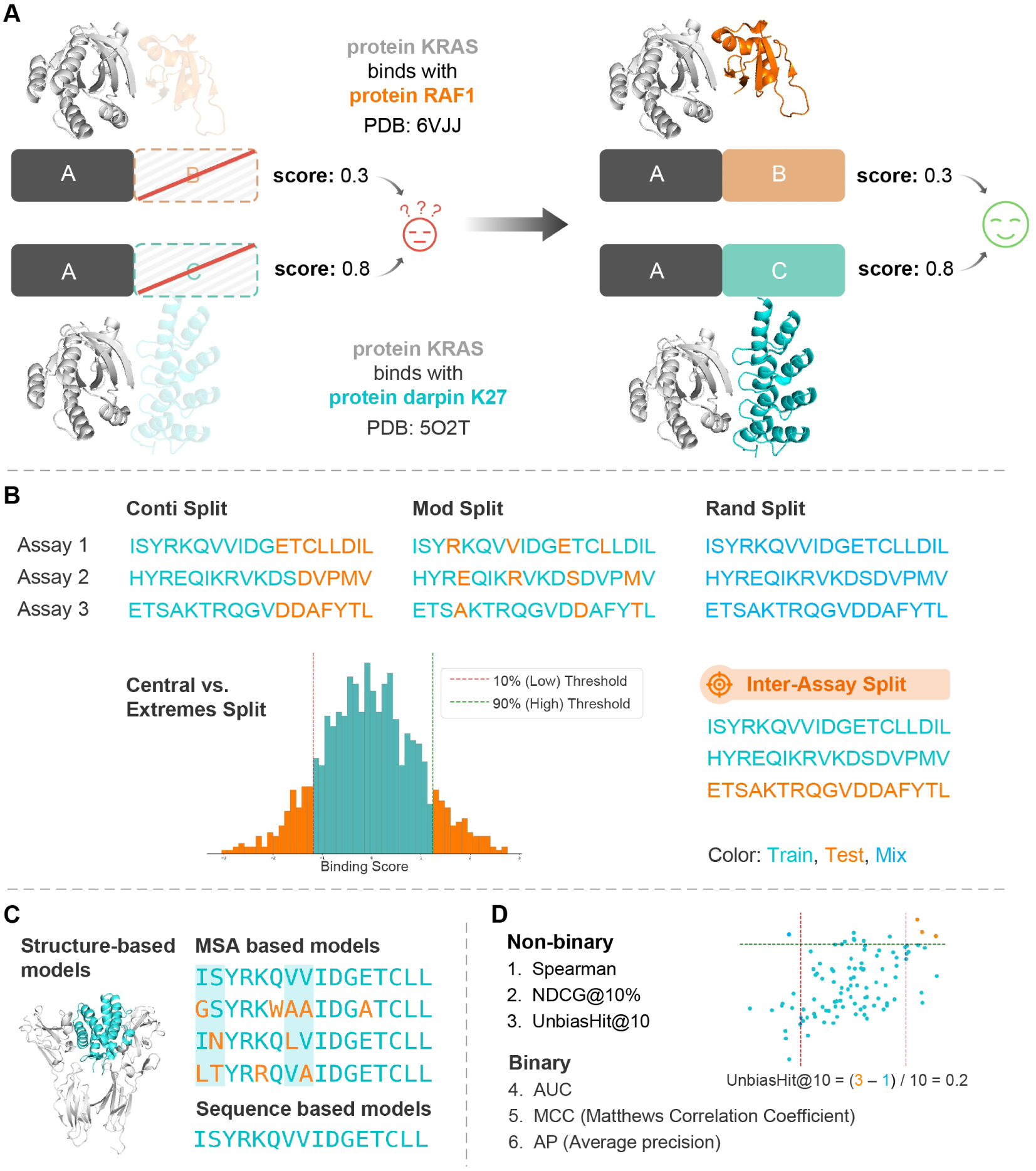
**A**, left: When the interacting partner is not modeled as in most datasets focusing on protein fitness, the same protein can have different binding scores across various assays, leading to confusion. Right: Modeling the full complex, as in BindingGYM, clarifies differences in binding scores, aiding the learning of the underlying physics of protein-protein interactions. **B**, We have implemented five different splits to examine model generalization capabilities, notably ’central vs. extremes split’ and ’inter-assay split’ to mimic real-world scenarios. **C**, Ten baseline models are included across three categories: structure-based, MSA-based, and sequence-based. **D**, Six evaluation metrics from two groups, binary and non-binary, are used. ’UnbiasHit@10’ measures the difference between the proportion of top ten scored mutants in the top 10% and those in the bottom 10%.

The fourth scheme, ’Central vs. Extremes Split’, sorts the entries by binding score, using the middle 80% for training, while the lowest and highest 10% form the test set. This approach aligns with the common optimization goal of using existing data to design variants with even higher scores. Including the lowest 10% helps reduce bias in evaluation and prevents the trained model from indiscriminately predicting high scores for new mutants.

The fifth, ’Inter-Assay Split’, is crucial, where a set of assays is used for training and a different set of assays for testing. This split evaluates the models’ ability to generalize to new assays, which holds significant practical significance.

### 3.3 Baseline models

We include ten baseline models across four categories. Language-based models are notably accessible, requiring only protein sequences as input. These include ProGen2, ESM1v, and ESM2 [45, 46, 47], which exclusively leverage protein sequence data.

Multi-sequence alignment (MSA)-based models such as EVE, Tranception, and TranceptEVE [48, 49, 50] extract evolutionary information from sequence databases. The latter two models also incorporate elements from protein language models, thereby enhancing their prediction capabilities.

Additionally, our dataset includes the structures of protein complexes, supporting the use of structure-based models such as ESM-IF1, ProteinMPNN, PPIformer, and SaProt [51, 52, 53, 54], which leverage these structures to predict protein-protein interactions. With rapid advancements in deep learning for structure prediction [55, 56, 23], the availability of structural data for protein monomers and complexes is expanding, enhancing the applicability of these structure-based models.

For each protein-protein pair, the score for each mutant is defined as the log-ratio of the probability of the mutant to the wild type, 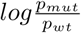 following [13, 57].

### 3.4 Metrics

We use six metrics to assess model performance: Spearman, AUC, MCC, NDCG, AP, and a specially designed metric called “UnbiasHit@10”. AUC (Area Under the ROC Curve) measures the model’s ability to discriminate between mutants with higher than binding affinity than the wild type and mutant with lower binding affinity. MCC (Matthews Correlation Coefficient) evaluates the quality of binary classifications, useful in imbalanced datasets. NDCG (Normalized Discounted Cumulative Gain) assesses the ranking quality of the predictions, valuing the order of relevance, we set the threshold at 10%, the same as [13]. AP (Average Precision) calculates the average precision value across different recall levels, highlighting precision-recall trade-offs. “UnbiasHit@10” is useful in practical scenarios where typically only about ten molecules undergo experimental testing with low-throughput, high-accuracy methods. This metric measures the difference between the proportion of the top 10 scored mutants that fall within the top 10% of actual performance and those that fall within the bottom 10%. This metric simulates the situation where we propose ten mutants with the highest predicted binding affinity for experimental validation.

## 4 Experiments

### 4.1 Evaluation of zero-shot performances

Due to the high costs associated with setting up experimental assays to quantitatively measure the binding energy between specific protein-protein pairs, zero-shot capability is crucial for discovering potential binders for a protein of interest (POI), studying the mutational effects on a POI, and designing novel binders to a target protein. In this section, we benchmark ten baseline models spanning three categories: structure-based, protein language based, and multiple sequence alignment (MSA) based. Unlike previous studies where predicted monomer structures from AlphaFold were used [13], we input full protein complex structures into our structure-based methods.

Table 2 indicates that models leveraging both evolutionary information from MSA and features from protein language models across a broader sequence database outperform those using either source alone. A structure-based method, ProteinMPNN, shows the best performance on our dataset. We anticipate that integrating evolutionary and physical interaction data more effectively could further enhance zero-shot performance.

**Table 2:** Zero-shot performance on predicting mutational effects on protein-protein interactions.

| Category | Model | Spearman | AUC | MCC | NDCG | AP | UnbiasHit@10 |
| --- | --- | --- | --- | --- | --- | --- | --- |
| Structure-based | ProteinMPNN | **0.40** | **0.69** | **0.15** | **0.72** | **0.22** | 0.30 |
|  | ESM-if1 | 0.34 | 0.66 | 0.14 | 0.70 | 0.20 | 0.22 |
|  | PiFold | 0.34 | 0.66 | 0.14 | 0.70 | 0.20 | 0.14 |
|  | ByProt | 0.28 | 0.62 | 0.10 | 0.67 | 0.17 | 0.02 |
|  | PPIformer | 0.19 | 0.61 | 0.06 | 0.59 | 0.14 | 0.04 |
|  | SaProt | 0.27 | 0.64 | 0.10 | 0.67 | 0.18 | 0.22 |
| Protein | ProGen2 | 0.25 | 0.61 | 0.09 | 0.66 | 0.16 | 0.14 |
| Language-based | ESM1v | 0.26 | 0.62 | 0.08 | 0.66 | 0.16 | 0.20 |
|  | ESM2 | 0.29 | 0.62 | 0.09 | 0.67 | 0.17 | 0.17 |
|  | ESM3 | 0.27 | 0.61 | 0.09 | 0.66 | 0.17 | 0.06 |
| MSA-based | EVE | 0.32 | 0.64 | 0.12 | 0.69 | 0.20 | 0.28 |
| MSA+Protein | Tranception | 0.32 | 0.65 | 0.12 | 0.69 | 0.20 | **0.31** |
| Language-based | TranceptEVE | 0.34 | 0.66 | 0.13 | 0.69 | 0.20 | 0.28 |

### 4.2 Evaluation of intra-assay finetuned performances

In protein optimization, where some experimental data for candidate protein molecules are available, fine-tuning is critical to enhance desirable properties, such as increased binding to target proteins or reduced binding to undesired targets. Due to space constraints, we present the results for two representative intra-assay split schemes here and provide additional results for two other splits in the supplementary materials.

For finetuning all baseline models, we employ learning-to-rank techniques [31, 32], ensuring uniformity across experiments by using the same batch size for each model, with every batch drawn exclusively from the same assay. Specifically, ESM2 and its randomly initialized variant, ESM2-R, are finetuned using LoRA [58] to mitigate overfitting. For further details, refer to the supplementary materials.

Table 3 shows the results for the random split, which is commonly used as a sanity check to verify that the dataset is informative and that the experimental results are learnable, not arbitrary. We randomly divided each assay’s data points into five folds. As demonstrated in Table 3, One-hot encoding (OHE) successfully learns from randomly split data. This confirms the reasonableness of the experimental outcomes and aligns with the established understanding that the majority of mutations do not significantly affect the binding score [30]. Once the effects of key mutations are learned, predicting outcomes for mutants with additional non-impacting mutations becomes straightforward.

**Table 3:** Performance of fine-tuned models on predicting mutational effects in protein-protein interactions, evaluated over five-fold random splits.

| Category | Model | Mutational Depth | Spearman | AUC | MCC | NDCG | AP |
| --- | --- | --- | --- | --- | --- | --- | --- |
| Structure-based | ProteinMPNN-R | ALL | 0.58 | 0.78 | 0.25 | 0.78 | 0.31 |
|  |  | <3 | 0.51 | 0.75 | 0.20 | 0.74 | 0.28 |
|  |  | >=3 | 0.54 | 0.80 | 0.31 | 0.80 | 0.38 |
|  | ProteinMPNN | ALL | 0.75 | 0.87 | 0.45 | **0.90** | 0.51 |
|  |  | <3 | 0.73 | 0.87 | 0.43 | 0.88 | 0.49 |
|  |  | >=3 | 0.63 | 0.85 | 0.45 | 0.88 | 0.50 |
| Protein Language-based | ESM2-R | ALL | 0.36 | 0.68 | 0.11 | 0.69 | 0.19 |
|  |  | <3 | 0.29 | 0.65 | 0.08 | 0.65 | 0.17 |
|  |  | >=3 | 0.33 | 0.69 | 0.14 | 0.71 | 0.23 |
|  | ESM2 | ALL | **0.76** | 0.88 | 0.45 | **0.90** | 0.53 |
|  |  | <3 | 0.74 | 0.88 | 0.45 | 0.89 | 0.52 |
|  |  | >=3 | 0.66 | 0.86 | 0.43 | 0.87 | 0.52 |
| OHE | OHE | ALL | **0.76** | **0.89** | **0.49** | **0.90** | **0.56** |
|  |  | <3 | 0.74 | 0.88 | 0.49 | 0.89 | 0.55 |
|  |  | >=3 | 0.66 | 0.87 | 0.45 | 0.88 | 0.52 |

Structure-based methods and protein language-based methods, such as ESM2, achieve similar performances to OHE, with a Spearman correlation coefficient of 0.76, which approaches the upper limit of what the quality of DMS data allows. Notably, when initialized with random weights, both ProteinMPNN and ESM2 perform worse than One-hot encoding (OHE), likely due to overfitting. Moreover, ESM2, with significantly more parameters, performs even worse.

The second split demonstrated here is the ’Contig Split’, where mutated residues are grouped into five contiguous segments in sequence space. We restrict the mutational depth to single, the same as ProteinGYM, to prevent information leak. This arrangement presents a significant challenge as there is no overlap in mutated residues between any two groups, forcing the model to learn transferable features. As shown in Table 4, One-hot encoding (OHE) fails to identify any transferable features. Pre-training proves to be advantageous; both ProteinMPNN and ESM2 significantly outperform their counterparts initialized with random weights.

**Table 4:** Performance of fine-tuned models on predicting mutational effects in protein-protein interactions, evaluated over five-fold contig splits.

| Category | Model | Mutational Depth | Spearman | AUC | MCC | NDCG | AP |
| --- | --- | --- | --- | --- | --- | --- | --- |
| Structure-based | ProteinMPNN-R | Single | 0.22 | 0.61 | 0.07 | 0.61 | 0.16 |
|  | ProteinMPNN | Single | **0.50** | **0.71** | **0.20** | **0.74** | **0.26** |
| Protein Language-based | ESM2-R | Single | 0.18 | 0.60 | 0.06 | 0.63 | 0.14 |
|  | ESM2 | Single | 0.46 | 0.67 | 0.12 | 0.71 | 0.18 |
| OHE | OHE | Single | -0.15 | 0.44 | 0.00 | 0.45 | 0.10 |

### 4.3 Evaluation of inter-assay finetuned performance

A key contribution of our work is the introduction of the inter-assay split, where assays are clustered based on the sequences of the mutated proteins into five distinct groups. Data from one group are used exclusively for testing, while data from the remaining four groups are used for training. This approach aims to evaluate the generalizability of models to unseen protein-protein pairs, thereby improving zero-shot performance in future PPI experiments.

We analyzed three levels of mutational depth: ALL, <3, and >=3, as presented in Table 5. Generally, mutants with more mutations are harder to predict than those with fewer mutations. One-hot encoding (OHE) struggles to transfer knowledge to unseen assays. Random initialization of weights for ProteinMPNN and ESM2 reduces performance, yet these models manage to learn some transferable features. ProteinMPNN achieves the best results, showing a slight improvement from zero-shot performance. Although the improvements are marginal, we anticipate that with more sophisticated model designs, improved quality filtering of existing data, and the continued accumulation of more data, as shown in protein stability prediction[59], we may observe a scaling law effect: as data volume increases, the performance of fine-tuned models also rises.

**Table 5:** Performance of fine-tuned models on predicting mutational effects in protein-protein interactions, evaluated over five-fold inter-assay splits.

| Category | Model | Mutational Depth | Spearman | AUC | MCC | NDCG | AP |
| --- | --- | --- | --- | --- | --- | --- | --- |
| Structure-based | ProteinMPNN-R | ALL | 0.16 | 0.57 | 0.05 | 0.59 | 0.14 |
|  |  | <3 | 0.11 | 0.56 | 0.04 | 0.56 | 0.15 |
|  |  | >=3 | 0.19 | 0.60 | 0.09 | 0.63 | 0.17 |
|  | ProteinMPNN | ALL | **0.42** | **0.70** | **0.16** | **0.72** | **0.23** |
|  |  | <3 | 0.43 | 0.70 | 0.16 | 0.72 | 0.22 |
|  |  | >=3 | 0.30 | 0.70 | 0.17 | 0.69 | 0.25 |
| Protein Language-based | ESM2-R | ALL | 0.09 | 0.55 | 0.03 | 0.57 | 0.13 |
|  |  | <3 | 0.09 | 0.55 | 0.02 | 0.56 | 0.13 |
|  |  | >=3 | 0.05 | 0.54 | 0.03 | 0.55 | 0.14 |
|  | ESM2 | ALL | 0.30 | 0.62 | 0.10 | 0.67 | 0.18 |
|  |  | <3 | 0.31 | 0.61 | 0.08 | 0.68 | 0.17 |
|  |  | >=3 | 0.15 | 0.60 | 0.08 | 0.60 | 0.18 |
| OHE | OHE | ALL | 0.00 | 0.50 | 0.00 | 0.00 | 0.10 |
|  |  | <3 | 0.00 | 0.50 | 0.00 | 0.00 | 0.10 |
|  |  | >=3 | 0.00 | 0.50 | 0.00 | 0.00 | 0.10 |

### 4.4 Comparison of model performance in zero-shot and finetuned setting

Using a five-fold inter-assay split allows us to generate predictions for all data, enabling a direct comparison between the performance of inter-assay fine-tuned models and their original zero-shot counterparts. As depicted in Fig 2, each dot represents an assay; in most cases, the fine-tuned models outperform the zero-shot models. For this analysis, we have set the performance benchmark of models initialized with random weights to zero in the zero-shot setting. ProteinMPNN demonstrates superior generalization compared to others.

**Figure 2:**
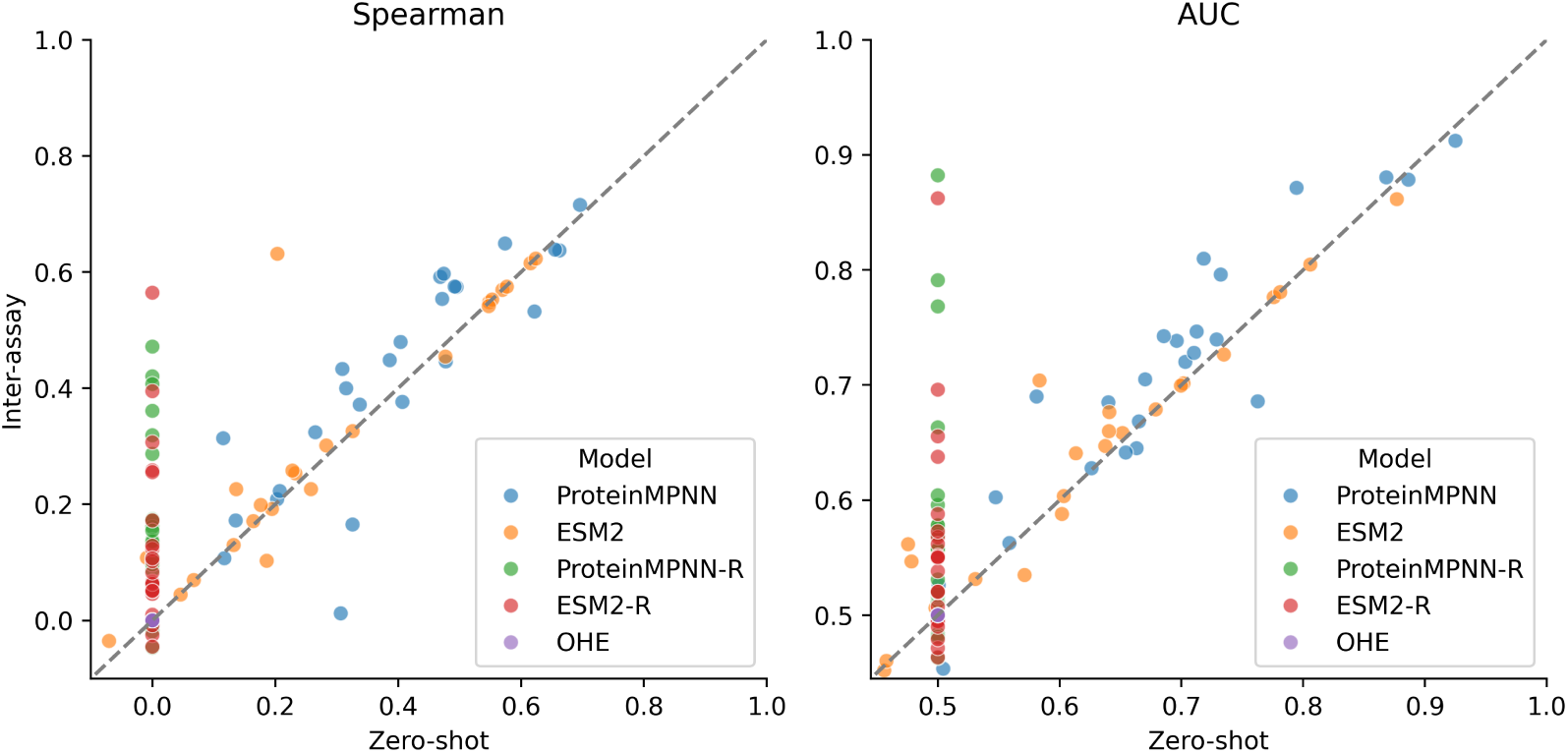
Each dot represents an assay. Finetuned models perform better than zero-shot, especially true for ProteinMPNN, where the dots lie above the diagonal line.

It is noteworthy that the two outlier points for ProteinMPNN, which fall below the diagonal line, are derived from the same study on protein co-evolution [60]. This suggests that these instances may involve more complex changes in interactions. For example, a mutation in protein A that is harmful in the original interaction environment might become benign or neutral when a reciprocal mutation occurs in protein B. Such dynamical change of protein structures underscore the complexity of protein-protein interactions and highlight the need for models that can adequately account for these protein dynamics.

## 5 Conclusions and Future Work

We have curated BindingGYM, the largest database of quantitative protein-protein interactions to date, highlighting the importance of modeling entire protein complexes. Each assay is meticulously paired with its corresponding complex structure. Five data split schemes were introduced, including two designed to simulate real-world scenarios: the ‘Central vs. Extremes Split’ for optimizing mutant binding and the ‘Inter-Assay Split’ for generalizing to new protein pairs. Our evaluation framework includes ten baseline models and six key metrics. Despite the strengths of structure-based models, which outperform sequence- or MSA-based methods and demonstrate superior generalization, there is still room for improvement. Current limitations include noise in DMS-generated data and the relatively limited number of protein-protein pairs studied. Nevertheless, structure-based approaches offer insights into transferable residue-level interactions from numerous mutations.

Looking ahead, as innovations in DMS experiments continue and next-generation sequencing becomes more affordable, the volume of available data will increase. We plan to update the BindingGYM database annually to ensure it remains comprehensive, setting the stage for a unified effort to decode the complex language of protein-protein interactions.

## Checklist

1. For all authors…

a. Do the main claims made in the abstract and introduction accurately reflect the paper’s contributions and scope? [Yes] see the experiments section.
b. Did you describe the limitations of your work? [Yes] See section Conclusions and Future Work.
c. Did you discuss any potential negative societal impacts of your work? [Yes] See section Supplementary material A.1.
d. Have you read the ethics review guidelines and ensured that your paper conforms to them? [Yes]
2. If you are including theoretical results…

a. Did you state the full set of assumptions of all theoretical results? [NA]
b. Did you include complete proofs of all theoretical results? [NA]
3. If you ran experiments (e.g. for benchmarks)…

a. Did you include the code, data, and instructions needed to reproduce the main experimental results (either in the supplemental material or as a URL)? [Yes] https://github.com/luwei0917/BindingGYM
b. Did you specify all the training details (e.g., data splits, hyperparameters, how they were chosen)? [Yes] in ’the BindingGYM dataset’ section and SI
c. Did you report error bars (e.g., with respect to the random seed after running experiments multiple times)? [Yes] In SI.
d. Did you include the total amount of compute and the type of resources used (e.g., type of GPUs, internal cluster, or cloud provider)? [Yes] In SI.
4. If you are using existing assets (e.g., code, data, models) or curating/releasing new assets…

a. If your work uses existing assets, did you cite the creators? [NA]
b. Did you mention the license of the assets? [NA]
c. Did you include any new assets either in the supplemental material or as a URL? [Yes] in SI.
d. Did you discuss whether and how consent was obtained from people whose data you’re using/curating? [Yes] in SI.
e. Did you discuss whether the data you are using/curating contains personally identifiable information or offensive content? [Yes] in SI.
5. If you used crowdsourcing or conducted research with human subjects…

a. Did you include the full text of instructions given to participants and screenshots, if applicable? [NA]
b. Did you describe any potential participant risks, with links to Institutional Review Board (IRB) approvals, if applicable? [NA]
c. Did you include the estimated hourly wage paid to participants and the total amount spent on participant compensation? [NA]

## A Appendix

### A.1 Broader societal impacts

Our research primarily focuses on the fundamental biophysical aspects of protein-protein interactions (PPI), which, by their nature, are not expected to directly result in negative societal impacts. While the development of improved PPI models could theoretically be applied in various contexts, including the design of proteins with potential biosecurity concerns, our study does not produce any new models specifically tailored for such applications. It is important, however, to acknowledge that any advancement in protein design technology carries potential dual-use concerns. Consequently, we advocate for responsible research and adherence to ethical standards to prevent misuse of scientific discoveries in this field.

### A.2 Detailed definitions of each column in Table 1

- **# of data**: Represents the total number of data points in the dataset, each corresponding to a unique entry derived from deep mutational scanning experiments, which capture variations in protein sequences and their respective binding energies.
- **HT (High Throughput)**: Indicates whether the data was generated using high-throughput techniques. A ‘Yes’ suggests that the dataset includes a large volume of data collected through automated processes, enabling comprehensive analysis at scale. A ‘No’ indicates traditional, lower-scale data collection methods.
- **C-Structure Available**: Specifies whether the crystal structure of the protein complex is available for the corresponding data entry. ‘Yes’ indicates that the complex structure data is available. ‘No’ means the complex structure is not available.
- **Quantitative**: Describes whether the data includes quantitative measurements of binding energies. ‘Yes’ indicates that the data provides numerical values representing binding affinities. ‘No’ suggests the data is qualitative or binary.
- **ML Ready**: Indicates if the data has been pre-processed and formatted to be directly used in machine learning models. ‘Yes’ means that the data is cleaned, normalized, and structured, making it immediately suitable for training predictive models. ‘No’ means that additional preprocessing might be required.
- **Multichain Support**: Indicates whether the dataset supports modeling of multiple protein chains. ‘Yes’ signifies that the data includes entries involving complex multichain interactions, which are essential for studying protein interactions. ‘No’ suggests that the dataset only supports modeling of single protein chains.
- **Design Usecase**: Describes the specific applications for which the dataset is designed, such as studying protein-protein interactions, protein networks, general protein fitness, protein representation learning, and therapeutic antibody design. This column highlights the potential research and development areas that can benefit from the dataset.

### A.3 Training Details

#### A.3.1 Data Partitioning

- **Intra_random**: Data is randomly divided into 5 folds using the KFold method from `sklearn.model_selection`, with a set seed of 42 to ensure reproducibility.
- **Intra_contig**: For assays with single-point mutation data count *≥* 100, the data is continuously segmented into 5 sections along the sequence, striving to ensure each segment has a similar amount of data.
- **Intra_mod**: For assays with single-point mutation data count *≥* 100, the data is divided into 5 segments based on the position modulo 5 (pos % 5).
- **Intra_two_extreme**: The bottom 10% and top 10% of the data are used as the test set, with the remaining data serving as the training set.
- **Inter_assay**: The target sequences are clustered using MMseqs2 [61] with a stringent sequence identity cutoff of 25%, to ensure meaningful generalization across different assays.

#### **A.3.2** Training

- **ProteinMPNN & ESM2**: All models are trained using the AdamW optimizer (learning rate = 0.001, weight decay = 0.05, epsilon = 0.00001) combined with the ListMLE loss [32]. Training proceeds for 100 epochs with an early stopping patience of 3 epochs. During training, all mutation positions are masked in the amino acids, and the predicted value is calculated as the difference between the sum of mutant logit probabilities and wild-type logit probabilities (∑(*mt_logit_probs*) − *∑*(*wt_logit_probs*))
- **OHE (One-Hot Encoding)**: Each sequence is transformed into a one-hot encoded feature matrix of dimensions (seq_len *×* 20), where seq_len is the length of the sequence and 20 represents the number of amino acid types. A Ridge regression model with an alpha value of 0.01 is then trained to directly predict the DMS scores, following methodologies similar to those described in [51].
- **OHE-AA (One-Hot Encoding of Amino Acid Mutations)**: Each sequence is encoded by comparing mutations to the wild-type (WT) sequence, forming a 20×20 matrix for one-hot encoding (e.g., a mutation from A to E at position 2 is encoded with ’1’ at the respective position in the matrix). This encoded data is then used to train a Ridge regression model with an alpha value of 0.01 to directly predict DMS scores.

### A.4 Total Amount of Compute and Type of Resources Used

The computational resources used for training and analysis comprised one NVIDIA A100 GPU, 48 CPUs, and 1024 GB of RAM.

### A.5 Ethical Considerations and Data Handling

- **Consent for Data Usage:** [Yes] Consent for using the data in the BindingGYM dataset was obtained through appropriate channels. The data primarily comprises publicly available deep mutational scanning (DMS) results, which are published in scientific literature. For any unpublished or privately sourced data, explicit consent was secured from the original contributors, ensuring compliance with ethical standards and data usage policies.
- **Personal Information and Offensive Content:** [No] The BindingGYM dataset does not include any personally identifiable information.

### A.6 Licensing Information

- **Usage License:** [Yes] The BindingGYM dataset is made available under the MIT License. This license permits reuse, distribution, and modification for both academic and commercial purposes, provided that proper credit is given to the original authors and the dataset. The MIT License is chosen for its permissiveness in encouraging open and collaborative scientific research, facilitating the widespread use and adaptation of the dataset in various biotechnological and pharmaceutical applications.

### A.7 Zero-shot performance with standard error bars

To assess the reliability and consistency of our zero-shot predictions, we computed standard error bars using a bootstrapping approach, as in Table 6. This method involved generating 1000 bootstrap samples from each assay to estimate the variability and confidence intervals around the predicted values.

**Table 6:** Zero-shot std error, computed based on 1000 bootstrap samples from the set of assays.

| Category | Model | Spearman | Spearman Std. error | AUC | AUC Std. error |
| --- | --- | --- | --- | --- | --- |
| Structure-based | ProteinMPNN | 0.40 | 0.03 | 0.69 | 0.02 |
|  | ESM-if1 | 0.34 | 0.04 | 0.66 | 0.02 |
|  | PiFold | 0.34 | 0.03 | 0.66 | 0.02 |
|  | ByProt | 0.28 | 0.04 | 0.62 | 0.02 |
|  | PPIformer | 0.19 | 0.04 | 0.61 | 0.02 |
|  | SaProt | 0.27 | 0.04 | 0.64 | 0.02 |
| Protein | ProGen2 | 0.25 | 0.04 | 0.61 | 0.02 |
| Language-based | ESM1v | 0.26 | 0.04 | 0.62 | 0.02 |
|  | ESM2 | 0.29 | 0.04 | 0.62 | 0.02 |
|  | ESM3 | 0.27 | 0.05 | 0.61 | 0.03 |
| MSA-based | EVE | 0.32 | 0.06 | 0.64 | 0.03 |
| MSA+Protein | Tranception | 0.32 | 0.04 | 0.65 | 0.02 |
| Language-based | TranceptEVE | 0.34 | 0.05 | 0.66 | 0.02 |

### A.8 Finetuned results for continuous split

Table 7 presents the finetuning results for baselines using the continuous split, where mutations within certain contiguous segments of the sequence space are designated for training, with the remaining segments used for testing. The effectiveness of the finetuned models is assessed on these test sets.

**Table 7:** Comparison of finetuning performance for continuous split.

| Category | Model | Mutational Depth | Spearman | AUC | MCC | NDCG | AP |
| --- | --- | --- | --- | --- | --- | --- | --- |
| Structure-based | ProteinMPNN-R | Single | 0.22 | 0.61 | 0.07 | 0.61 | 0.16 |
|  | ProteinMPNN | Single | 0.50 | 0.71 | 0.20 | 0.74 | 0.26 |
| Protein | ESM2-R | Single | 0.18 | 0.60 | 0.06 | 0.63 | 0.14 |
| Language-based | ESM2 | Single | 0.46 | 0.67 | 0.12 | 0.71 | 0.18 |
| OHE | OHE | Single | -0.15 | 0.44 | 0.00 | 0.45 | 0.10 |
|  | OHE-AA | Single | 0.12 | 0.55 | 0.02 | 0.59 | 0.13 |

ProteinMPNN displayed the best finetuned results, underlining the advantage of pre-training; conversely, ProteinMPNN with randomly initialized weights showed inferior performance. One-hot encoding, including OHE-AA, performed poorly, demonstrating limited ability to generalize to previously unseen regions.

### A.9 Finetuned results for modulo split

Table 8 demonstrates that one-hot encoding fails to generalize under the modulo split, while both ESM2 and ProteinMPNN achieve similar performance. This similarity likely arises because adjacent amino acids significantly influence the mutational effects on any specific amino acid.

**Table 8:**
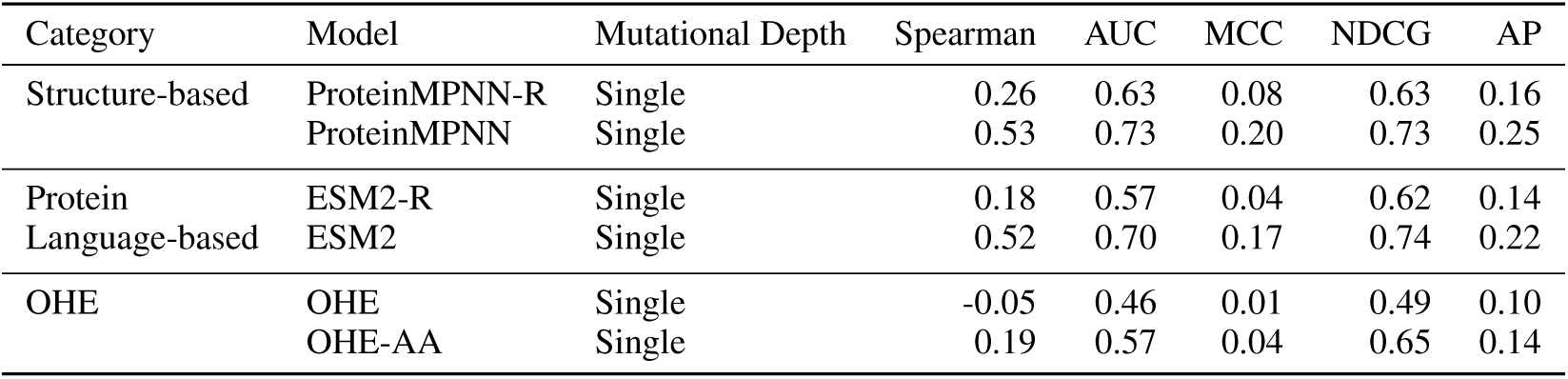
Comparison of finetuning performance for modulo split.

| Category | Model | Mutational Depth | Spearman | AUC | MCC | NDCG | AP |
| --- | --- | --- | --- | --- | --- | --- | --- |
| Structure-based | ProteinMPNN-R | Single | 0.26 | 0.63 | 0.08 | 0.63 | 0.16 |
|  | ProteinMPNN | Single | 0.53 | 0.73 | 0.20 | 0.73 | 0.25 |
| Protein<br>Language-based | ESM2-R | Single | 0.18 | 0.57 | 0.04 | 0.62 | 0.14 |
|  | ESM2 | Single | 0.52 | 0.70 | 0.17 | 0.74 | 0.22 |
| OHE | OHE | Single | -0.05 | 0.46 | 0.01 | 0.49 | 0.10 |
|  | OHE-AA | Single | 0.19 | 0.57 | 0.04 | 0.65 | 0.14 |

### A.10 Finetuned results for central vs extremes split

Table 9 illustrates that all baselines provide reasonable predictions in the Central vs Extremes split, where the strongest and weakest binders form the test set. This effectiveness largely stems from the nature of DMS assays, where the test set consists of mutants at the two extremes, often with multiple mutations. Since individual mutations present in test set mutants are likely already encountered in the training set, models are effectively able to identify mutants with improved binding affinities. But as shown in Table 7, 8 and 5 in the main text, generalizing to unseen positions or new assays is still very challenging.

**Table 9:** Comparison of finetuning performance for central vs extremes split.

| Category | Model | TopHit@10 | BottomHit@10 | UnbiasHit@10 |
| --- | --- | --- | --- | --- |
| Structure-based | ProteinMPNN-R | 0.54 | 0.13 | 0.41 |
|  | ProteinMPNN | 0.82 | 0.02 | 0.80 |
| Protein<br>Language-based | ESM2-R | 0.55 | 0.10 | 0.45 |
|  | ESM2 | 0.80 | 0.02 | 0.78 |
| OHE | OHE | 0.73 | 0.02 | 0.71 |

### A.11 Distribution of DMS score by assay

In Figure 3, we present the histograms showing the distribution of DMS scores for each assay included in the benchmark. These distributions are crucial for understanding the variability captured by different assays. The experimental setups vary significantly among assays, leading to notable differences in score distributions. In certain cases, the histogram peaks sharply at a specific value, indicating a high frequency of scores around that point, which may suggest the score for the wild type. Conversely, other assays exhibit bimodal distributions, where two distinct peaks suggest the presence of two different predominant groups. Moreover, the range of scores varies across assays, reflecting differences in the magnitude of mutational impacts or in the sensitivity of the assays. To accommodate these differences effectively and improve the predictive accuracy of our models, we employ ranking-based machine learning techniques, such as learning to rank. This approach allows us to handle the heterogeneity between assays and learn meaningful interactions from the relative rankings of mutations within each specific assay context.

**Figure 3:**
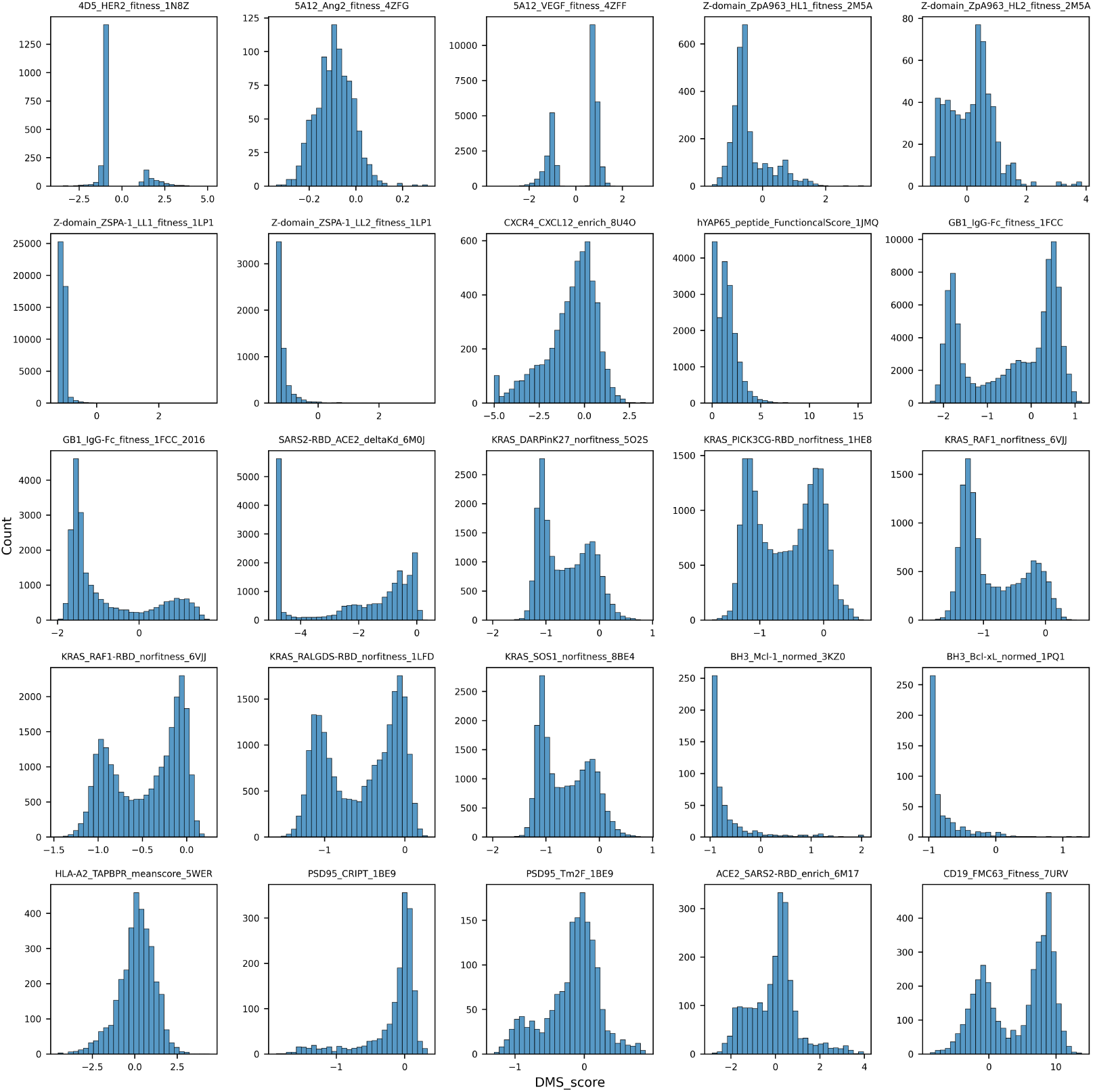
DMS score distribution.

### A.12 Pairwise target sequence similarity

Some assays involve the same types of proteins, such as KRAS and its binding partners, leading to shared similarities among entries. While a model trained on one such assay may perform well on others, our primary interest lies in the model’s ability to generalize to new targets. Therefore, we cluster target sequences and assess the inter-assay fine-tuned results exclusively across these clusters. Figure 4 illustrates the pairwise sequence similarity among all assays, highlighting potential overlaps and distinctions.

**Figure 4:**
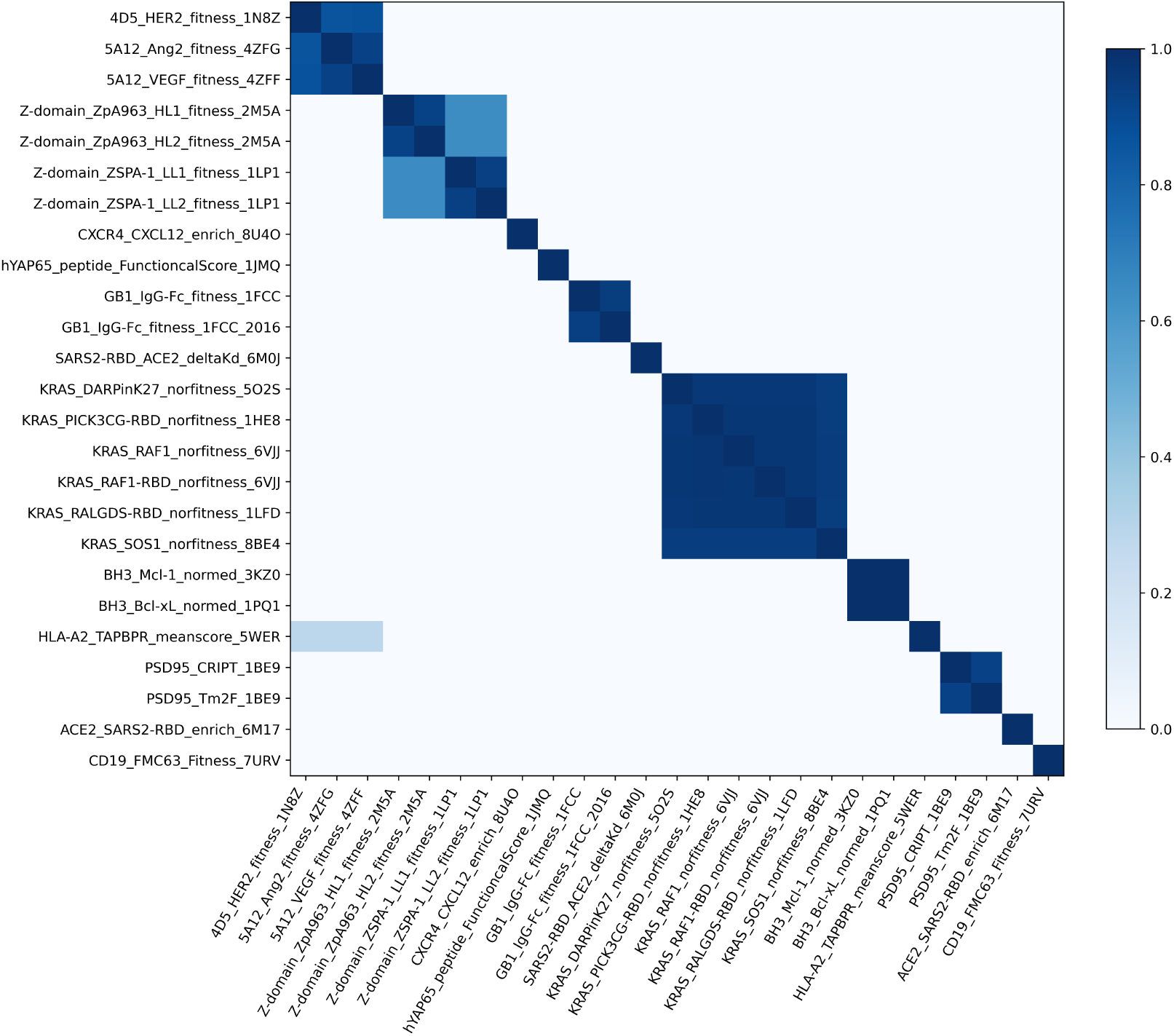
Target sequence similarity across all assays.

### A.13 Source and Target Amino Acids

Figure 5 displays the frequency of each specific amino acid being mutated to every other specific amino acid.

**Figure 5:**
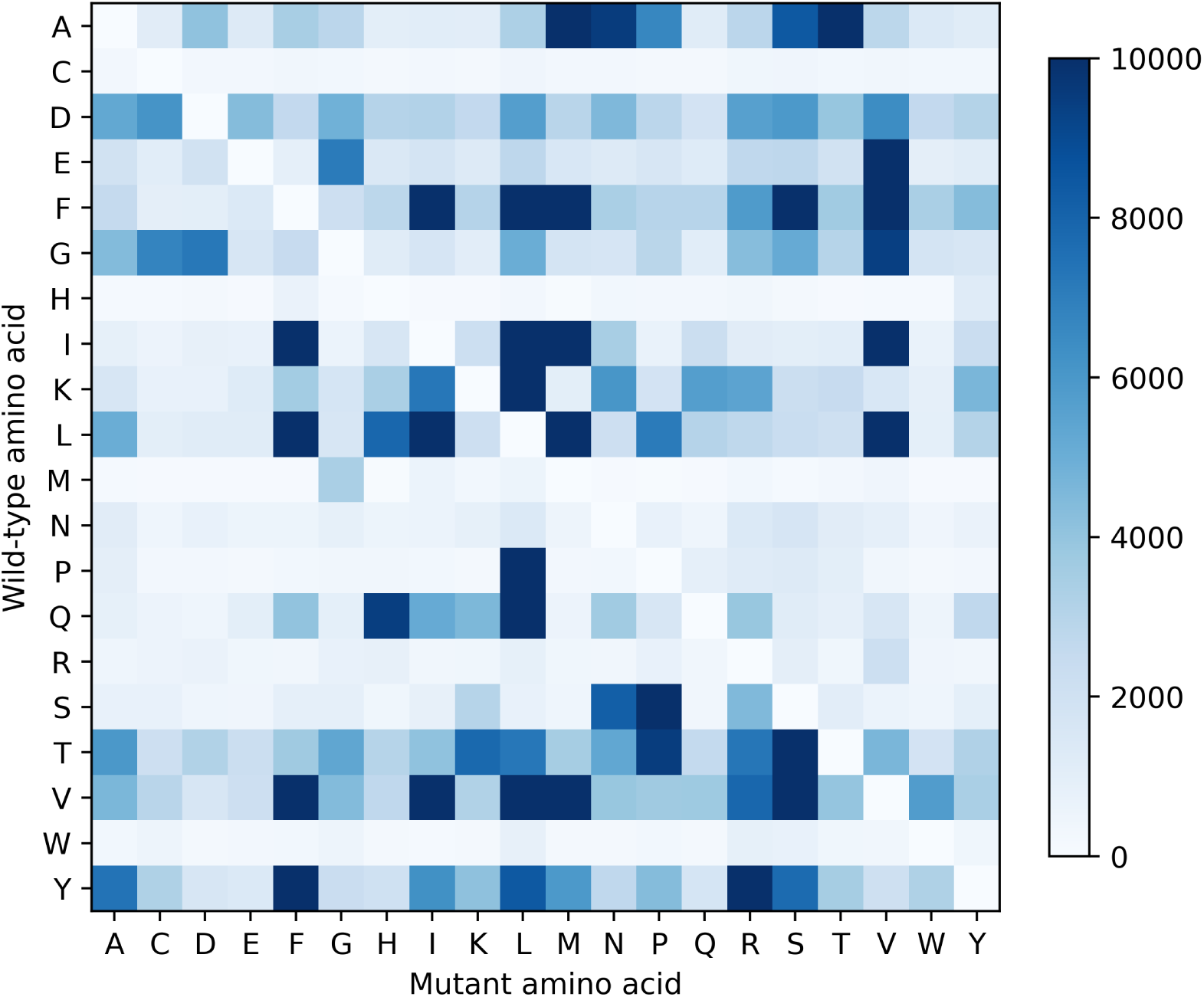
Distribution of amino acid mutations.

### A.14 List of data sources

Various studies on protein functions and interactions have been compiled, gathering diverse protein-protein interaction data, as in Table 10. These studies cover a wide range of protein types, including immunoglobulins, chemokine receptors, cytokine receptors, growth factors, regulatory proteins, and virus-related proteins. Additionally, the data includes interactions of the same protein with different other proteins, helping models understand different interaction regions of the same protein and enhancing the model’s generalization ability.

**Table 10:**
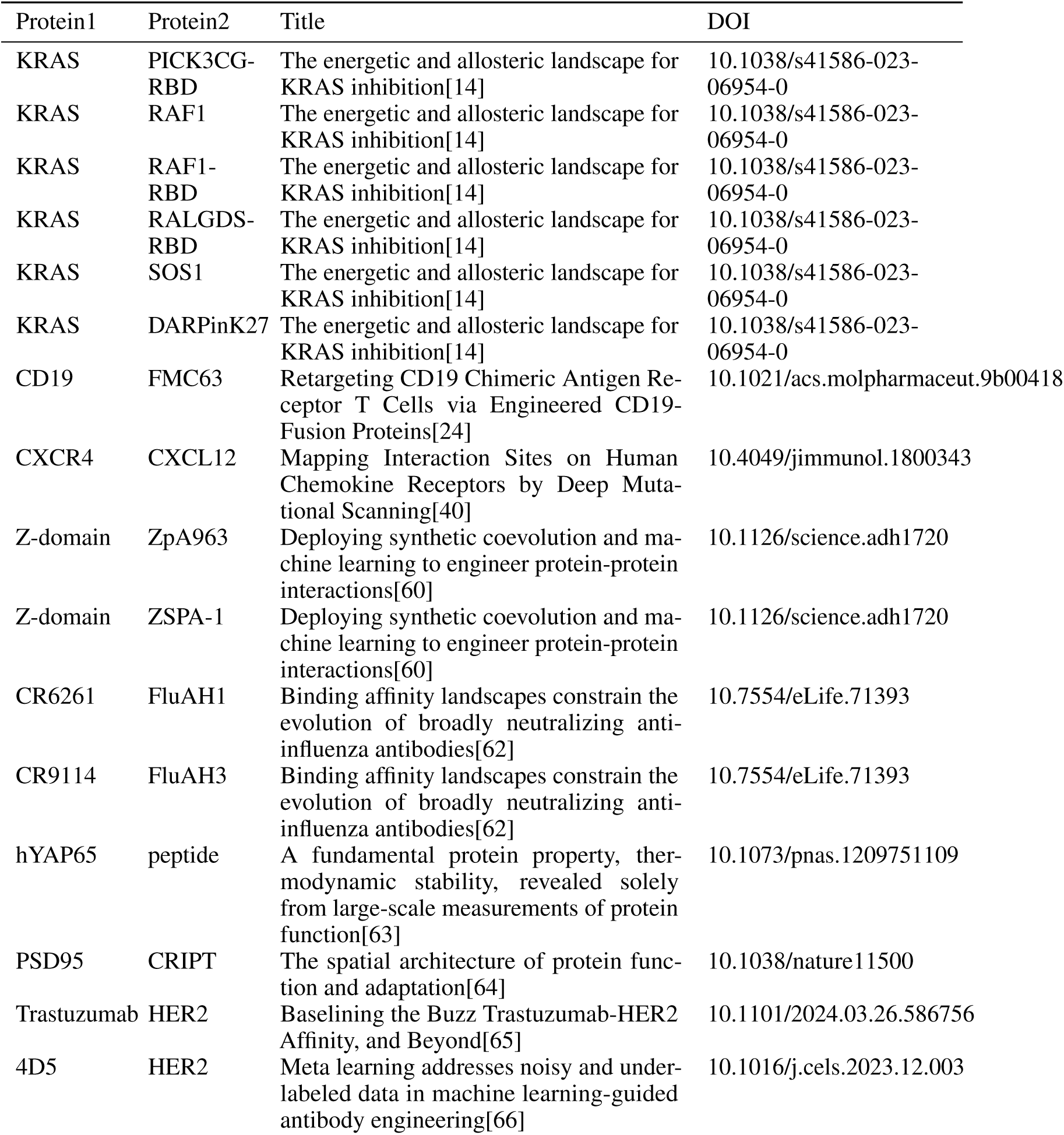
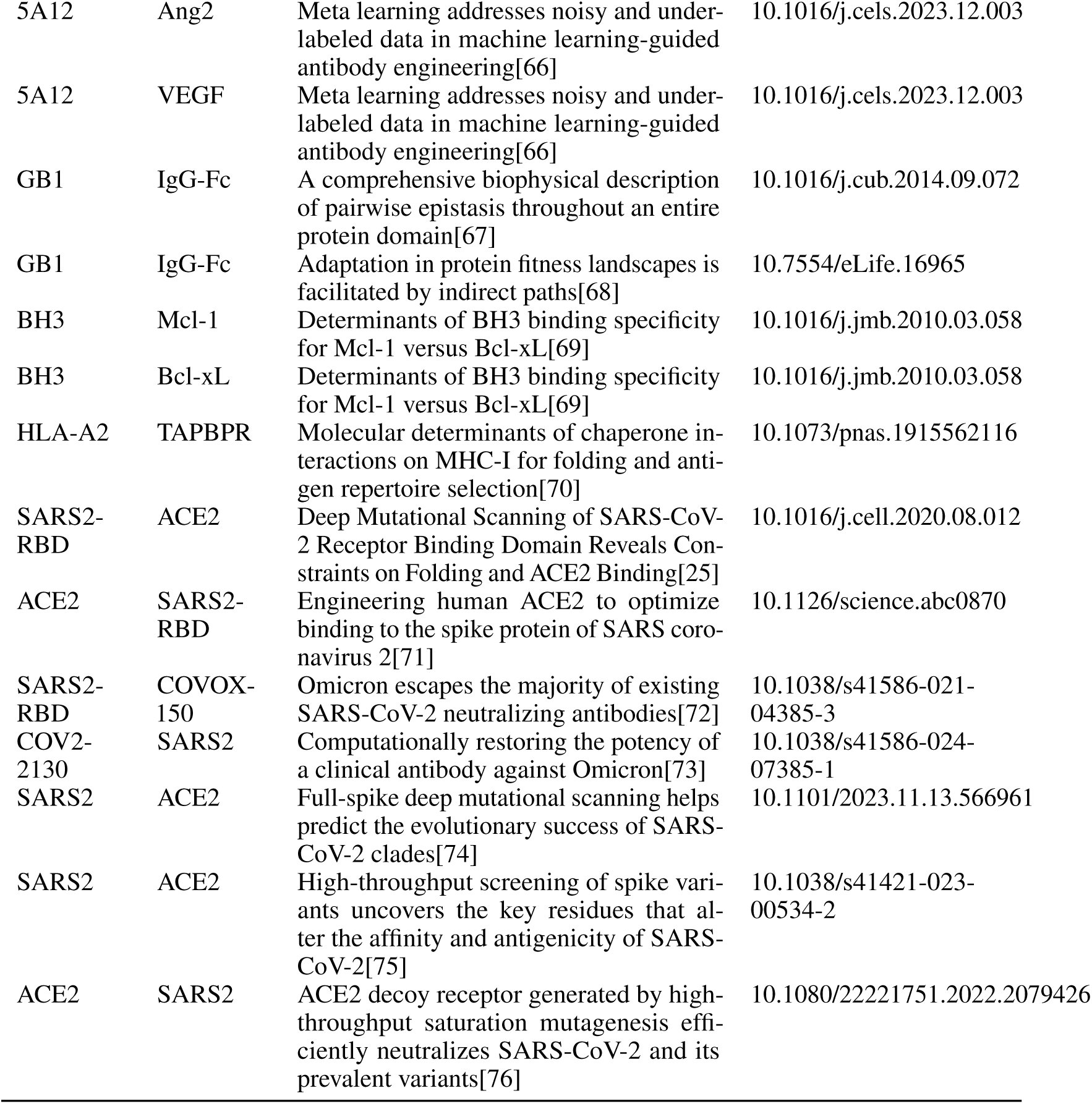
List of data sources.

| Protein1 | Protein2 | Title | DOI |
| --- | --- | --- | --- |
| KRAS | PICK3CG-RBD | The energetic and allosteric landscape for KRAS inhibition[14] | 10.1038/s41586-023-06954-0 |
| KRAS | RAF1 | The energetic and allosteric landscape for KRAS inhibition[14] | 10.1038/s41586-023-06954-0 |
| KRAS | RAF1-RBD | The energetic and allosteric landscape for KRAS inhibition[14] | 10.1038/s41586-023-06954-0 |
| KRAS | RALGDS-RBD | The energetic and allosteric landscape for KRAS inhibition[14] | 10.1038/s41586-023-06954-0 |
| KRAS | SOS1 | The energetic and allosteric landscape for KRAS inhibition[14] | 10.1038/s41586-023-06954-0 |
| KRAS | DARPinK27 | The energetic and allosteric landscape for KRAS inhibition[14] | 10.1038/s41586-023-06954-0 |
| CD19 | FMC63 | Retargeting CD19 Chimeric Antigen Receptor T Cells via Engineered CD19-Fusion Proteins[24] | 10.1021/acs.molpharmaceut.9b00418 |
| CXCR4 | CXCL12 | Mapping Interaction Sites on Human Chemokine Receptors by Deep Mutational Scanning[40] | 10.4049/jimmunol.1800343 |
| Z-domain | ZpA963 | Deploying synthetic coevolution and machine learning to engineer protein-protein interactions[60] | 10.1126/science.adh1720 |
| Z-domain | ZSPA-1 | Deploying synthetic coevolution and machine learning to engineer protein-protein interactions[60] | 10.1126/science.adh1720 |
| CR6261 | FluAH1 | Binding affinity landscapes constrain the evolution of broadly neutralizing anti-influenza antibodies[62] | 10.7554/eLife.71393 |
| CR9114 | FluAH3 | Binding affinity landscapes constrain the evolution of broadly neutralizing anti-influenza antibodies[62] | 10.7554/eLife.71393 |
| hYAP65 | peptide | A fundamental protein property, thermodynamic stability, revealed solely from large-scale measurements of protein function[63] | 10.1073/pnas.1209751109 |
| PSD95 | CRIPT | The spatial architecture of protein function and adaptation[64] | 10.1038/nature11500 |
| Trastuzumab | HER2 | Baselining the Buzz Trastuzumab-HER2 Affinity, and Beyond[65] | 10.1101/2024.03.26.586756 |
| 4D5 | HER2 | Meta learning addresses noisy and under-labeled data in machine learning-guided antibody engineering[66] | 10.1016/j.cels.2023.12.003 |
| 5A12 | Ang2 | Meta learning addresses noisy and under-labeled data in machine learning-guided antibody engineering[66] | 10.1016/j.cels.2023.12.003 |
| 5A12 | VEGF | Meta learning addresses noisy and under-labeled data in machine learning-guided antibody engineering[66] | 10.1016/j.cels.2023.12.003 |
| GB1 | IgG-Fc | A comprehensive biophysical description of pairwise epistasis throughout an entire protein domain[67] | 10.1016/j.cub.2014.09.072 |
| GB1 | IgG-Fc | Adaptation in protein fitness landscapes is facilitated by indirect paths[68] | 10.7554/eLife.16965 |
| BH3 | Mcl-1 | Determinants of BH3 binding specificity for Mcl-1 versus Bcl-xL[69] | 10.1016/j.jmb.2010.03.058 |
| BH3 | Bcl-xL | Determinants of BH3 binding specificity for Mcl-1 versus Bcl-xL[69] | 10.1016/j.jmb.2010.03.058 |
| HLA-A2 | TAPBPR | Molecular determinants of chaperone interactions on MHC-I for folding and antigen repertoire selection[70] | 10.1073/pnas.1915562116 |
| SARS2-RBD | ACE2 | Deep Mutational Scanning of SARS-CoV-2 Receptor Binding Domain Reveals Constraints on Folding and ACE2 Binding[25] | 10.1016/j.cell.2020.08.012 |
| ACE2 | SARS2-RBD | Engineering human ACE2 to optimize binding to the spike protein of SARS coronavirus 2[71] | 10.1126/science.abc0870 |
| SARS2-RBD | COVOX-150 | Omicron escapes the majority of existing SARS-CoV-2 neutralizing antibodies[72] | 10.1038/s41586-021-04385-3 |
| COV2-2130 | SARS2 | Computationally restoring the potency of a clinical antibody against Omicron[73] | 10.1038/s41586-024-07385-1 |
| SARS2 | ACE2 | Full-spike deep mutational scanning helps predict the evolutionary success of SARS-CoV-2 clades[74] | 10.1101/2023.11.13.566961 |
| SARS2 | ACE2 | High-throughput screening of spike variants uncovers the key residues that alter the affinity and antigenicity of SARS-CoV-2[75] | 10.1038/s41421-023-00534-2 |
| ACE2 | SARS2 | ACE2 decoy receptor generated by high-throughput saturation mutagenesis efficiently neutralizes SARS-CoV-2 and its prevalent variants[76] | 10.1080/22221751.2022.2079426 |

**Table 11:** Zero-shot Performance of models with error bar.

| Category | Model | Spearman | AUC | MCC | NDCG | AP | UnbiasHit@10 |
| --- | --- | --- | --- | --- | --- | --- | --- |
| Structure-based | ProteinMPNN | 0.40 ± 0.03 | 0.69 ± 0.02 | 0.15 ± 0.03 | 0.72 ± 0.02 | 0.22 ± 0.03 | 0.30 ± 0.05 |
| | ESM-if1 | 0.34 ± 0.04 | 0.66 ± 0.02 | 0.14 ± 0.03 | 0.70 ± 0.03 | 0.20 ± 0.02 | 0.22 ± 0.06 |
| | PiFold | 0.34 ± 0.04 | 0.66 ± 0.02 | 0.14 ± 0.03 | 0.70 ± 0.02 | 0.20 ± 0.02 | 0.14 ± 0.06 |
| | PPIformer | 0.19 ± 0.04 | 0.61 ± 0.02 | 0.06 ± 0.01 | 0.59 ± 0.02 | 0.14 ± 0.01 | 0.04 ± 0.05 |
| | SaProt | 0.27 ± 0.04 | 0.64 ± 0.02 | 0.10 ± 0.02 | 0.67 ± 0.03 | 0.18 ± 0.02 | 0.22 ± 0.05 |
| Protein<br>Language-based | ProGen2 | 0.25 ± 0.04 | 0.61 ± 0.02 | 0.09 ± 0.02 | 0.66 ± 0.03 | 0.16 ± 0.02 | 0.14 ± 0.06 |
| | ESM1v | 0.26 ± 0.04 | 0.62 ± 0.02 | 0.08 ± 0.02 | 0.66 ± 0.03 | 0.16 ± 0.02 | 0.20 ± 0.06 |
| | ESM2 | 0.29 ± 0.04 | 0.62 ± 0.02 | 0.09 ± 0.03 | 0.67 ± 0.03 | 0.17 ± 0.02 | 0.17 ± 0.06 |
| | ESM3 | 0.27 ± 0.05 | 0.61 ± 0.03 | 0.09 ± 0.03 | 0.66 ± 0.03 | 0.17 ± 0.02 | 0.06 ± 0.06 |
| MSA-based | EVE | 0.32 ± 0.06 | 0.64 ± 0.03 | 0.12 ± 0.03 | 0.69 ± 0.03 | 0.20 ± 0.02 | 0.28 ± 0.06 |
| MSA+Protein<br>Language-based | Tranception | 0.32 ± 0.04 | 0.65 ± 0.02 | 0.12 ± 0.03 | 0.69 ± 0.03 | 0.20 ± 0.02 | 0.31 ± 0.07 |
| | TranceptEVE | 0.34 ± 0.05 | 0.66 ± 0.02 | 0.13 ± 0.03 | 0.69 ± 0.03 | 0.20 ± 0.02 | 0.28 ± 0.05 |

**Table 12:** Performance of models with error bar, evaluated over five-fold random splits.

| Category | Model | Mutational Depth | Spearman | AUC | MCC | NDCG | AP |
| --- | --- | --- | --- | --- | --- | --- | --- |
| Structure-based | ProteinMPNN-R | ALL | 0.58 ± 0.04 | 0.78 ± 0.02 | 0.25 ± 0.03 | 0.78 ± 0.02 | 0.31 ± 0.03 |
| | | <3 | 0.51 ± 0.04 | 0.75 ± 0.02 | 0.20 ± 0.03 | 0.74 ± 0.02 | 0.28 ± 0.02 |
| | | >=3 | 0.54 ± 0.07 | 0.80 ± 0.04 | 0.31 ± 0.06 | 0.80 ± 0.04 | 0.38 ± 0.06 |
| | ProteinMPNN | ALL | 0.75 ± 0.04 | 0.87 ± 0.02 | 0.45 ± 0.03 | 0.90 ± 0.01 | 0.51 ± 0.03 |
| | | <3 | 0.73 ± 0.04 | 0.87 ± 0.02 | 0.43 ± 0.03 | 0.88 ± 0.02 | 0.49 ± 0.03 |
| | | >=3 | 0.63 ± 0.06 | 0.85 ± 0.04 | 0.45 ± 0.06 | 0.88 ± 0.03 | 0.50 ± 0.07 |
| Protein<br>Language-based | ESM2-R | ALL | 0.36 ± 0.03 | 0.68 ± 0.02 | 0.11 ± 0.02 | 0.69 ± 0.02 | 0.19 ± 0.02 |
| | | <3 | 0.29 ± 0.03 | 0.65 ± 0.02 | 0.08 ± 0.02 | 0.65 ± 0.02 | 0.17 ± 0.01 |
| | | >=3 | 0.33 ± 0.05 | 0.69 ± 0.03 | 0.14 ± 0.04 | 0.71 ± 0.04 | 0.23 ± 0.03 |
| | ESM2 | ALL | 0.76 ± 0.04 | 0.88 ± 0.02 | 0.45 ± 0.03 | 0.90 ± 0.02 | 0.53 ± 0.04 |
| | | <3 | 0.74 ± 0.04 | 0.88 ± 0.02 | 0.45 ± 0.03 | 0.89 ± 0.02 | 0.52 ± 0.03 |
| | | >=3 | 0.66 ± 0.06 | 0.86 ± 0.04 | 0.43 ± 0.06 | 0.87 ± 0.03 | 0.52 ± 0.07 |
| OHE | OHE | ALL | 0.76 ± 0.03 | 0.89 ± 0.02 | 0.49 ± 0.04 | 0.90 ± 0.02 | 0.56 ± 0.04 |
| | | <3 | 0.74 ± 0.04 | 0.88 ± 0.02 | 0.49 ± 0.04 | 0.89 ± 0.02 | 0.55 ± 0.04 |
| | | >=3 | 0.66 ± 0.05 | 0.87 ± 0.03 | 0.45 ± 0.06 | 0.88 ± 0.03 | 0.52 ± 0.07 |

**Table 13:** Performance of models with error bar, evaluated over five-fold contig splits.

| Category | Model | Mutational Depth | Spearman | AUC | MCC | NDCG | AP |
| --- | --- | --- | --- | --- | --- | --- | --- |
| Structure-based | ProteinMPNN-R | Single | $0.22 \pm 0.03$ | $0.61 \pm 0.02$ | $0.07 \pm 0.02$ | $0.61 \pm 0.02$ | $0.16 \pm 0.02$ |
| | ProteinMPNN | Single | $0.50 \pm 0.04$ | $0.71 \pm 0.03$ | $0.20 \pm 0.03$ | $0.74 \pm 0.02$ | $0.26 \pm 0.03$ |
| Protein<br>Language-based | ESM2-R | Single | $0.18 \pm 0.02$ | $0.60 \pm 0.01$ | $0.06 \pm 0.02$ | $0.63 \pm 0.02$ | $0.14 \pm 0.01$ |
| | ESM2 | Single | $0.46 \pm 0.04$ | $0.67 \pm 0.02$ | $0.12 \pm 0.03$ | $0.71 \pm 0.02$ | $0.18 \pm 0.02$ |
| OHE | OHE | Single | $-0.15 \pm 0.04$ | $0.44 \pm 0.03$ | $0.00 \pm 0.01$ | $0.45 \pm 0.03$ | $0.10 \pm 0.01$ |

**Table 14:**
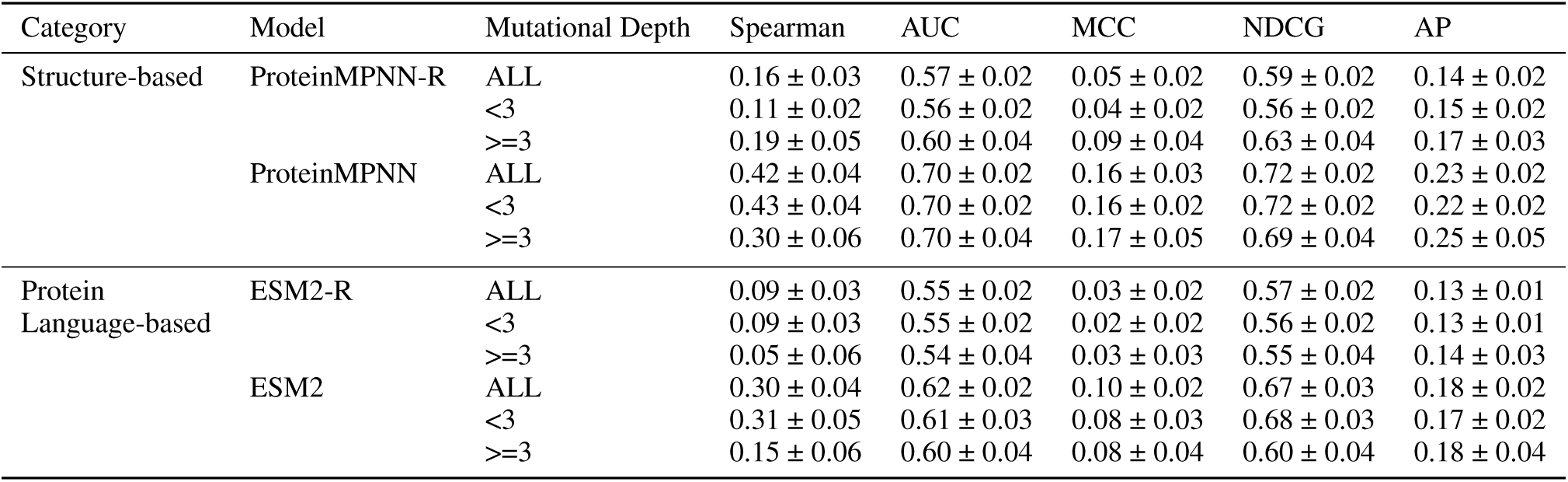
Performance of models with error bar, evaluated over five-fold inter-assay splits.

| Category | Model | Mutational Depth | Spearman | AUC | MCC | NDCG | AP |
| --- | --- | --- | --- | --- | --- | --- | --- |
| Structure-based | ProteinMPNN-R | ALL | $0.16 \pm 0.03$ | $0.57 \pm 0.02$ | $0.05 \pm 0.02$ | $0.59 \pm 0.02$ | $0.14 \pm 0.02$ |
| | | <3 | $0.11 \pm 0.02$ | $0.56 \pm 0.02$ | $0.04 \pm 0.02$ | $0.56 \pm 0.02$ | $0.15 \pm 0.02$ |
| | | >=3 | $0.19 \pm 0.05$ | $0.60 \pm 0.04$ | $0.09 \pm 0.04$ | $0.63 \pm 0.04$ | $0.17 \pm 0.03$ |
| | ProteinMPNN | ALL | $0.42 \pm 0.04$ | $0.70 \pm 0.02$ | $0.16 \pm 0.03$ | $0.72 \pm 0.02$ | $0.23 \pm 0.02$ |
| | | <3 | $0.43 \pm 0.04$ | $0.70 \pm 0.02$ | $0.16 \pm 0.02$ | $0.72 \pm 0.02$ | $0.22 \pm 0.02$ |
| | | >=3 | $0.30 \pm 0.06$ | $0.70 \pm 0.04$ | $0.17 \pm 0.05$ | $0.69 \pm 0.04$ | $0.25 \pm 0.05$ |
| Protein<br>Language-based | ESM2-R | ALL | $0.09 \pm 0.03$ | $0.55 \pm 0.02$ | $0.03 \pm 0.02$ | $0.57 \pm 0.02$ | $0.13 \pm 0.01$ |
| | | <3 | $0.09 \pm 0.03$ | $0.55 \pm 0.02$ | $0.02 \pm 0.02$ | $0.56 \pm 0.02$ | $0.13 \pm 0.01$ |
| | | >=3 | $0.05 \pm 0.06$ | $0.54 \pm 0.04$ | $0.03 \pm 0.03$ | $0.55 \pm 0.04$ | $0.14 \pm 0.03$ |
| | ESM2 | ALL | $0.30 \pm 0.04$ | $0.62 \pm 0.02$ | $0.10 \pm 0.02$ | $0.67 \pm 0.03$ | $0.18 \pm 0.02$ |
| | | <3 | $0.31 \pm 0.05$ | $0.61 \pm 0.03$ | $0.08 \pm 0.03$ | $0.68 \pm 0.03$ | $0.17 \pm 0.02$ |
| | | >=3 | $0.15 \pm 0.06$ | $0.60 \pm 0.04$ | $0.08 \pm 0.04$ | $0.60 \pm 0.04$ | $0.18 \pm 0.04$ |

